# Mechanosensitive remodeling of the bacterial flagellar motor is independent of direction of rotation

**DOI:** 10.1101/2021.01.19.427295

**Authors:** Navish Wadhwa, Yuhai Tu, Howard C. Berg

## Abstract

Motility is critical for the survival and dispersal of bacteria, and it plays an important role during infection. How bacteria regulate motility is thus a question of broad interest. Regulation of bacterial motility by chemical stimuli is well studied, but recent work has added a new dimension to the problem of motility control. The bidirectional flagellar motor of the bacterium *Escherichia coli* recruits or releases torque-generating units (stator units) in response to changes in load. Here, we show that this mechanosensitive remodeling of the flagellar motor is independent of direction of rotation. Remodeling rate constants in clockwise rotating motors and in counterclockwise rotating motors, measured previously, fall on the same curve if plotted against torque. Increased torque decreases the off rate of stator units from the motor, thereby increasing the number of active stator units at steady state. A simple mathematical model based on observed dynamics provides quantitative insight into the underlying molecular interactions. The torque-dependent remodeling mechanism represents a robust strategy to quickly regulate output (torque) in response to changes in demand (load).

**Significance:** Macromolecular machines carry out most of the biological functions in living organisms. Despite their significance, we do not yet understand the rules that govern the self-assembly of large multi-protein complexes. The bacterial flagellar motor tunes the assembly of its torque-generating stator complex with changes in external load. Here, we report that clockwise and counterclockwise rotating motors have identical remodeling response to changes in the external load, suggesting a purely mechanical mechanism for this regulation. Autonomous control of self-assembly may be a general strategy for tuning the functional output of protein complexes. The flagellar motor is a prime example of a macromolecular machine in which the functional regulation of assembly can be rigorously studied.

## Introduction

Motility is critical for many bacteria as it enables resource acquisition, the dispersal of progeny, and infection (1, 2). The rotation of flagella (3, 4), powered by the bidirectional flagellar motor (5–7), drives motility in many bacteria. In *Escherichia coli*, the flagellar motor consists of over 20 different proteins that self-assemble at the cell wall in varying copy numbers (8–10). Motor structure (Fig. 1A) includes a rotor embedded in the inner cell membrane, a drive shaft, and a flexible hook that transmits torque to the filament (10, 11). The C-ring, which contains copies of the proteins FliG, FliM, and FliN, is mounted on the cytoplasmic face of the rotor and is responsible for directional switching of the motor (12). The rotor is driven by up to 11 ion-powered MotA_5_B_2_ stator units (13–16) that surround the rotor and generate torque. MotA engages FliG whereas MotB is mounted on the rigid framework of the peptidoglycan cell wall (17–20). Motor-bound units exchange with a pool of unbound units in the inner membrane (10, 21).

**Fig. 1.**
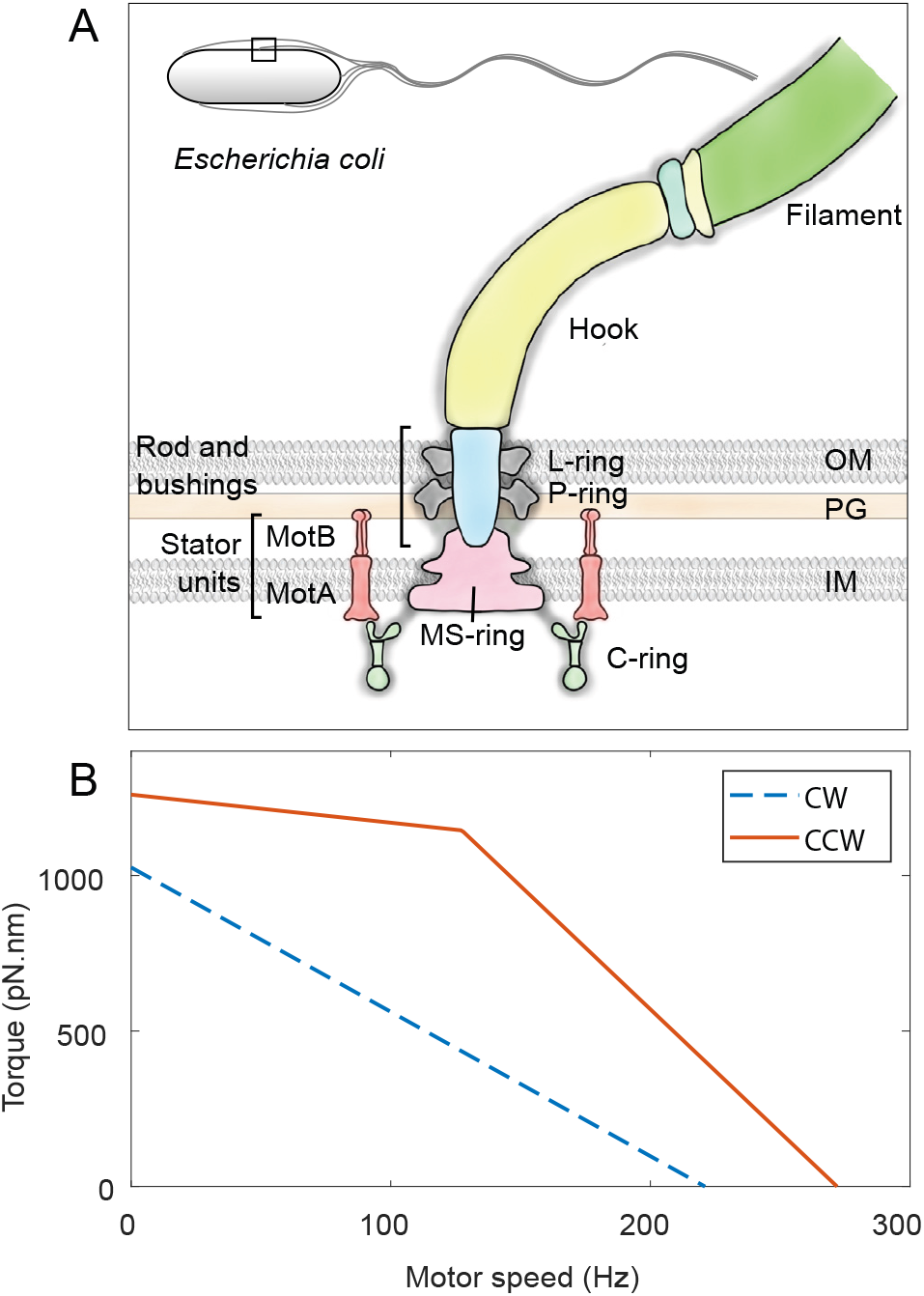
Bacterial flagellar motor’s structure and its torque-speed curve. **A**. Schematic representation of the flagellar motor of Gram-negative bacteria. The rotor consists of the MS-ring embedded in the inner membrane (IM) and the C-ring embedded in the cytoplasm. Stator units (MotA-MotB complexes) that span the inner membrane bind to the peptidoglycan (PG) layer and apply torque on the C-ring. The torque is transmitted via a rod (driveshaft) and a flexible hook (universal joint) to the flagellar filament. L- and P-rings are embedded in the outer membrane (OM) and the peptidoglycan (PG), respectively, and act as bushings. Inset shows the outline of an *E. coli* cell with a square demarcating the region that is represented in detail. **B**. Torque-speed curve of the counterclockwise (solid orange) and clockwise (dashed blue) rotating flagellar motors compared in this study. Adapted from refs. (28, 31). See *Materials and Methods* for details.

Motor function is regulated by inputs from the environment. Detection of specific ligands by chemoreceptors drives a two-component signaling cascade that controls the direction of rotation of the motor (22–24). Upon binding the response regulator CheY-P, the C-ring undergoes a concerted conformational change that reverses motor rotation from counter-clockwise (CCW) to clockwise (CW), as viewed from outside the cell. This change in the direction of rotation is the basis of run-and-tumble motility in *E. coli* (CCW = run, CW = tumble). Changes in viscous load trigger remodeling of the stator (25–27), whereby at high loads, the number of motor-bound stator units increases, and *vice versa*. Dynamics of stator remodeling have only been quantified in CCW rotating motors, using electrorotation (28) and magnetic tweezers (29, 30). The observed dynamics were rationalized using the CCW torque-speed relationship (Fig. 1B) (28). CCW and CW rotating motors have different torque-speed relationships (Fig. 1B), likely due to differences in stator-rotor interactions (31–33). How the differences in torque-speed relationship affect stator remodeling in CW motors is unknown. Additionally, the molecular mechanisms underlying the load-dependent remodeling phenomenon remain poorly understood.

Here, we use electrorotation to study the dynamics of load-dependent stator remodeling in CW rotating motors. We find that just like CCW motors, CW rotating flagellar motors release their stator units when the motor torque is low, and recruit stator units when the torque increases again. Remarkably, the rates of stator unit release and recruitment in CW and CCW motors collapse onto a single curve when plotted against torque, despite their dissimilar torque-speed relationships. The collapse of remodeling data suggests a universal model for torque dependence in the mechanically regulated remodeling of the bacterial flagellar motor. Our *in vivo* measurements of stator assembly dynamics advance the understanding of a large protein complex with multiple parts.

## Results

CCW motors have a unique torque-speed relationship among rotary molecular motors; the torque exerted by CCW motors remains nearly constant at low speeds up to a so-called “knee-speed”, beyond which it rapidly drops to zero (Fig. 1B). In contrast, torque produced by CW rotating motors drops linearly from stall to zero (31). In addition, the CW mutant strain used here has a significantly smaller stall torque and zero-torque speed than that of the CCW mutant strain used for comparison (Fig. 1B; also see Methods). These differences allow us to tease apart the roles of motor torque and speed in load-dependent stator remodeling.

We tethered bacterial cells to the surface of a sapphire window via short filament stubs (Fig. 2A). With the filament immobilized, the motor rotated the cell body at a low speed and operated close to stall. We observed motor output in this manner for 30 s. Then we turned on the electrorotation field, which applied an assisting external torque on the cell (see Methods), thereby decreasing the load on the motor. As a result, the cell rotation sped up and the motor torque decreased. To observe any changes in the number of active stator units, we turned electrorotation OFF for 1 s every 9 s, when the motor rotation due solely to the bound stator units could be measured. We repeated this cycle of 8 s ON followed by 1 s OFF for 10 min, after which we kept electrorotation off. Removal of electrorotation field removed the assisting torque and increased the load on the motor. The speed of the motor was now measured continuously. We observed the motor rotation in this manner for an additional 10 min.

**Fig. 2.**
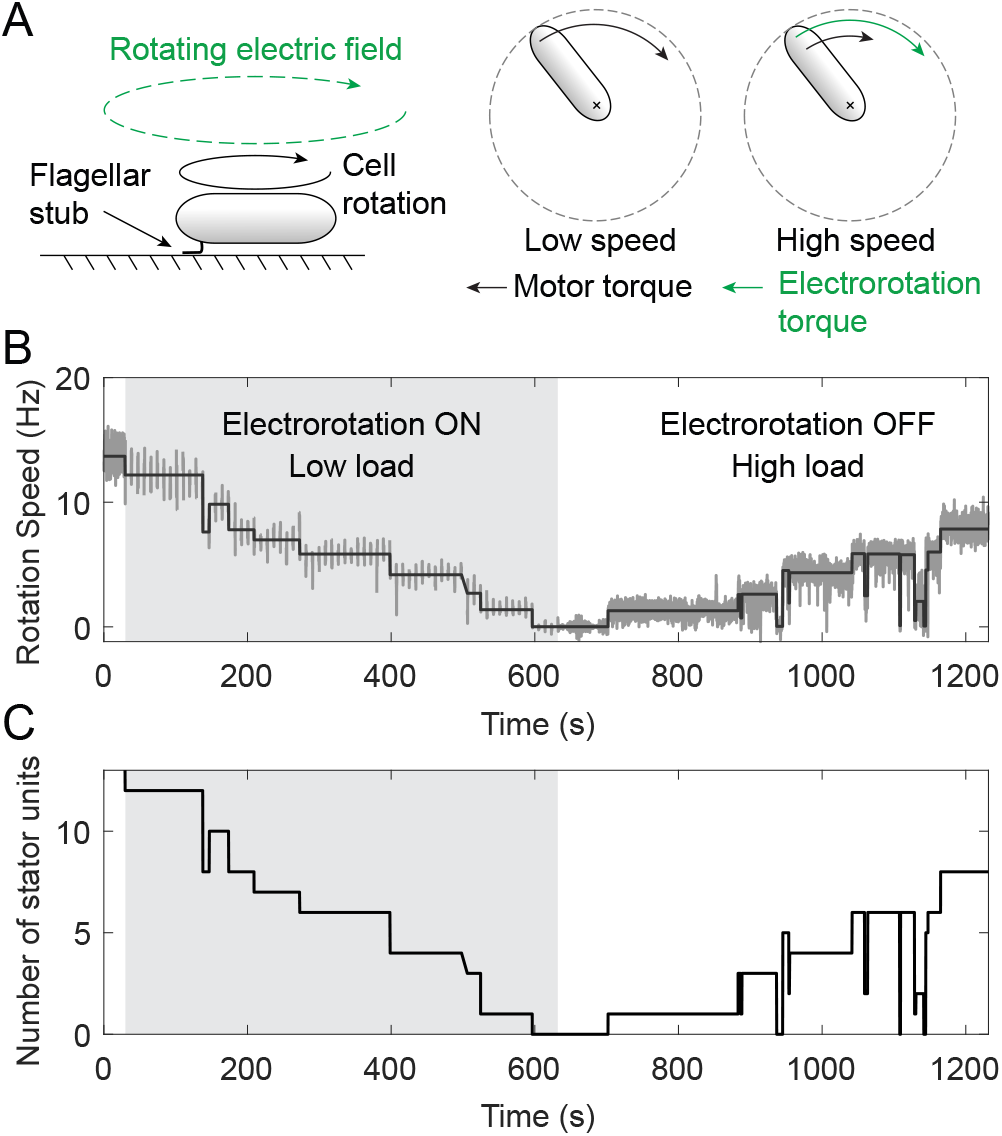
Control of motor load using electrorotation. **A**. The cell is tethered to a surface via a short flagellar stub (left). The motor exerts a large torque to rotate the cell body, depicted by the black arrow (middle). A high-frequency rotating electric field applies an external torque (green) on the cell. The external torque spins the cell at high speed and reduces the motor torque (right). Also see Methods. **B**. Example of an experimental output showing motor speed (dark gray) as a function of time for an electrorotation speed of 200 Hz. At the start of the experiment, motor speed was measured continuously for 30 s without any stimuli, after which electrorotation was turned on, depicted by the light gray region. During electrorotation, motor speed was measured for 1 s every 9 s. After 10 min of electrorotation, the field was turned off, increasing the load on the motor. Solid black line represents steps fitted on the speed data. **C**. Number of active stator units vs. time for the same data.

When we decreased motor load by driving a tethered cell forward with electrorotation, the motor’s native speed (measured during the OFF intervals) decreased, indicating a loss of stator units driving the motor. Fig. 2B shows the results of a typical experiment, in which the electrorotation speed during the ON intervals was fixed at 200 Hz. Starting at ∼14 Hz before electrorotation, this motor’s speed decreased in a stepwise manner to 0 Hz during the electrorotation period, indicating a complete loss of the bound stator units. The removal of electrorotation field after 10 min promoted the recruitment of stator units, indicated by a stepwise increase in the motor speed. We fitted steps to the speed data, from which we estimated the unitary step height corresponding to the gain or loss of single stator units. By dividing the speed levels by the unitary step height, we calculated the number of bound stator units as a function of time (Fig. 2C).

We conducted these experiments at five electrorotation speeds, ranging from 50 Hz to 250 Hz. These speeds cover the entire range of torque generation by the CW rotating motors, which decreases linearly from stall torque at 0 Hz to zero torque at 221 Hz (Fig. 1B). The dynamics of stator remodeling depended strongly on the electrorotation speed. We observed no remodeling for electrorotation at 50 Hz within the duration of the experiment (Fig. 3A). For the other electrorotation speeds, we observed a clear response to the reduction in load. Higher electrorotation speeds resulted in a bigger loss of stator units and at a greater rate (Fig. 3B-E). In all cases, the loss of the stator units during the electrorotation period was followed by a period of recovery after electrorotation was switched off.

**Fig. 3.**
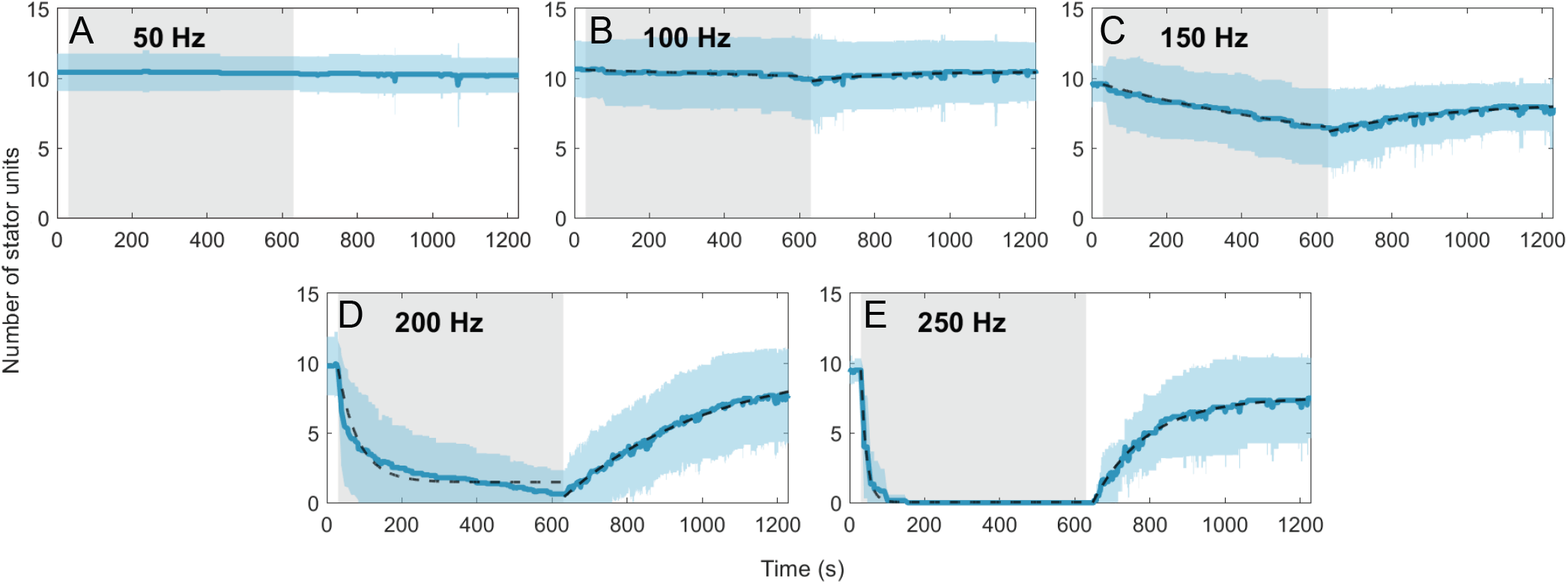
Number of stator units vs. time for different CW electrorotation speeds. The solid blue line is the average for a given electrorotation speed, and the shaded blue region represents the SD. Fits of Hill-Langmuir model (Eq. 2) on the data are depicted as dashed lines, except in panel A (electrorotation at 50 Hz) in which no remodeling was observed. The shaded gray region represents the electrorotation period. Sample sizes for A-E are 14, 15, 18, 19, and 6, respectively.

The kinetics of stator remodeling can be described by the Hill-Langmuir model as adopted by Nord *et al*. (29). An unbound stator unit can occupy one of the available sites on the periphery of the rotor with *N* binding sites. We fitted this model to the measured dynamics (Fig. 3), obtaining the on rate *k*_+_ and the off rate *k*_−_ for the interaction between a single stator unit and the motor. Most of the variation between experimental conditions was contained in *k*_−_ (Fig. 4A). In contrast, *k*_+_ measured during the electrorotation period showed little variation across experimental conditions (Fig. 4B); *k*_+_ was, however, higher during the recovery period in which the torque was high. We compared these rates with the data we obtained for CCW rotating motors (28). Remarkably, the two sets of data collapse if plotted against the torque per stator unit Γ (Fig. 4A). No such collapse was seen when the data were plotted against the motor rotation speed (Fig. 4A, B inset). This is striking because CW and CCW rotating motors produce different torques at any given speed (Fig. 1B) and have different remodeling kinetics at a given electrorotation speed. The collapse of the two data sets despite these differences demonstrates that torque is the main parameter governing load-dependent stator remodeling.

**Fig. 4.**
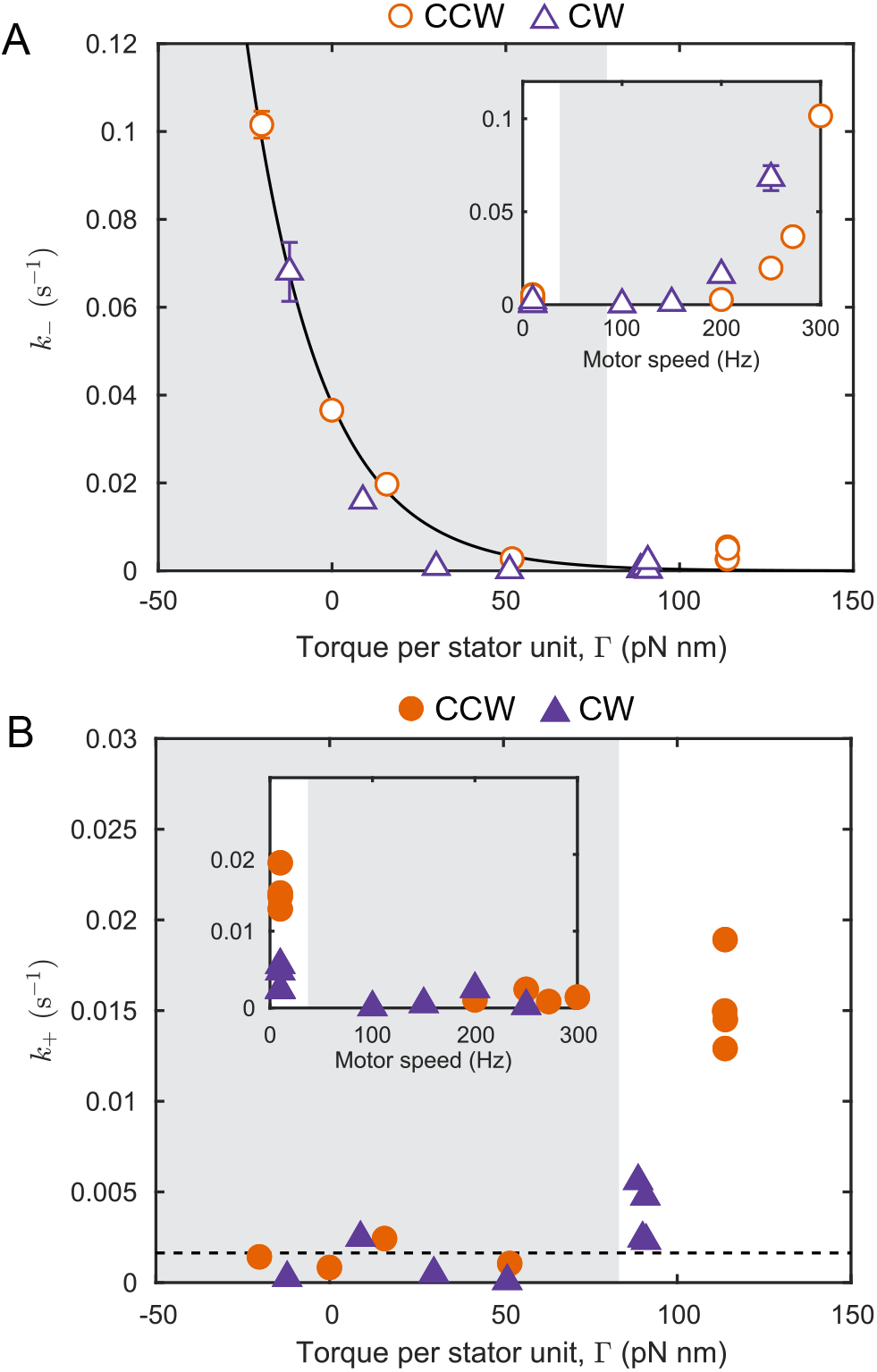
Data collapse if plotted against torque. **A**. The off rate *k_–_* (open symbols) as a function of torque per stator unit Γ, for CW (purple triangles) and CCW (orange disks) rotating motors. The solid line is the model 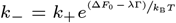, with *k*_+_ = 0.0016 s^−1^, Δ*F*_0_ = 3.2 *k*_B_*T*, and *λ* = 0.047 *k*_B_*T* .pN^−^ .nm. *k*_B_ and *T* are the Boltzmann constant and the absolute temperature, respectively. Inset shows the same data plotted against motor rotation speed. **B**. The on rate *k*_+_ (solid symbols) as a function of Γ, for CW (purple triangles) and CCW (orange disks) rotating motors. The dashed line is *k*_+_ = 0.0016 s^−1^. Inset shows the same data plotted against motor rotation speed.

The difference between the effective free energy of bound and unbound stator units (Δ*F*) can be defined from logarithm of the ratio of the forward and backward rates 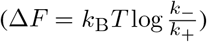, and determines whether the binding is favorable (Fig. 5A). Plotted against torque per stator unit (Γ), the values of Δ*F* measured for CW and CCW motors also collapsed onto a single curve, which exhibits a linear decrease in Δ*F* with Γ. The slope of a linear fit was given by *λ* = 0.047 *k*_B_*T* .pN^−1^.nm^−1^. But note that *λ* is a dimensionless quantity (energy and torque have the same units), giving *λ* = 0.19 in dimensionless units. The intercept of the linear fit was positive at Δ*F*_0_ ≈ 3.2 *k*_B_*T*, indicating that at zero-torque, the binding of stator units to the motor is unfavorable. A model of *k*_−_ based on the linear fit of Δ*F* against Γ captured the observed dependence of *k*_−_ on torque (Fig. 4A).

**Fig. 5.**
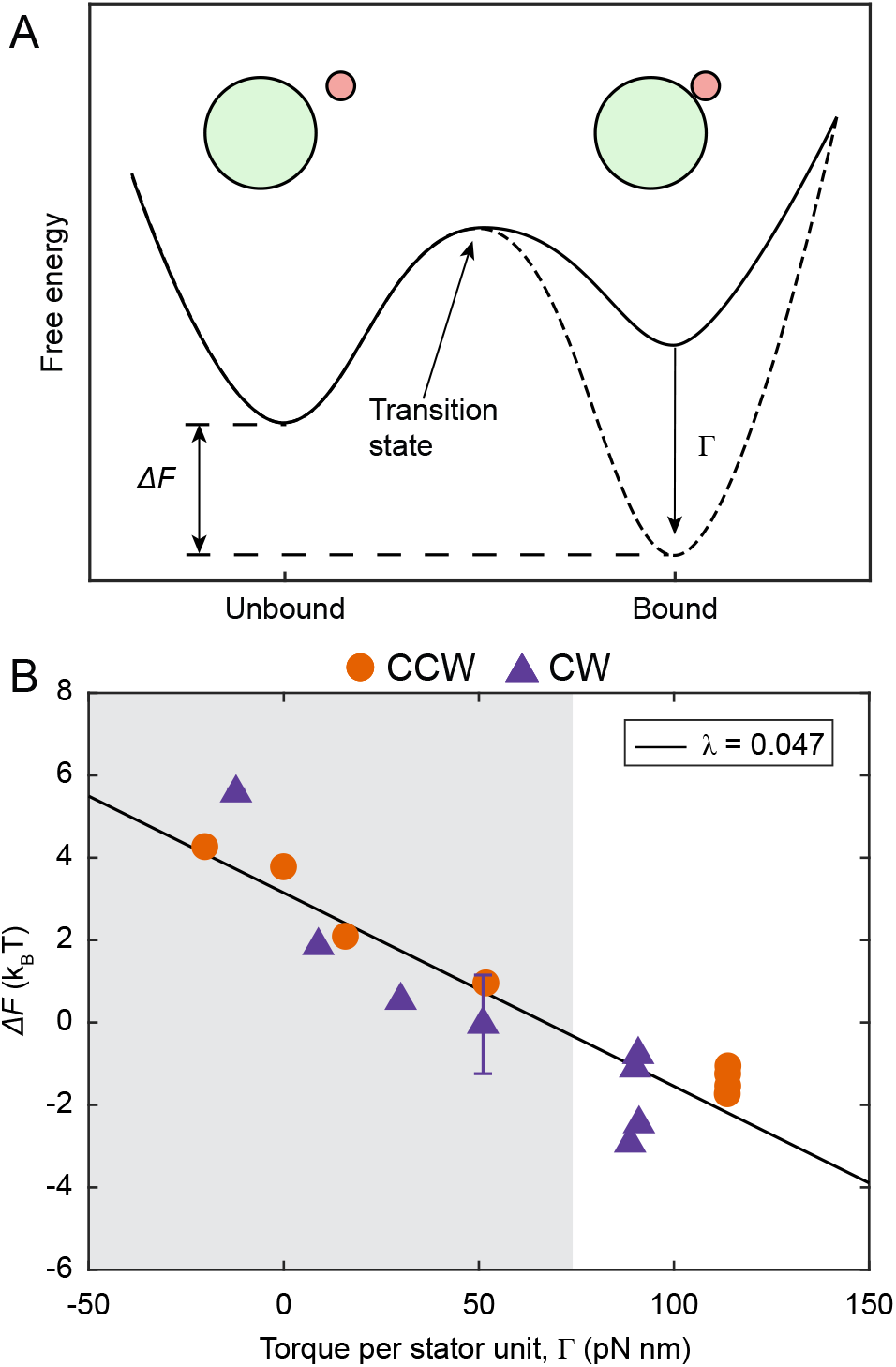
Torque tunes the effective free energy profile of stator binding. **A**. Schematic representation of how increased torque promotes stator assembly. At low torque (solid curve), the free energy of the bound stator units is greater than that of un-bound units, making assembly unfavorable. Increase in torque reduces the free energy of bound units (dashed curve), making assembly favorable. Δ*F* is the difference between the free energy of bound and unbound units. **B**. Δ*F* as a function of torque per stator unit Γ, for CW (purple triangles) and CCW (orange disks) rotating motors. The solid black line is linear fit to the data, and has a slope *λ* = 0.047 *k*_B_*T* .pN^−1^ .nm^−1^ or 0.19 in dimensionless units. The shaded and blank regions indicate data obtained from the electrorotation and the recovery periods, respectively. Error bars are 1 SE in each direction.

## Discussion

Our results show not only that load-dependent stator remodeling takes place in flagellar motors rotating in either direction, but that the remodeling dynamics as a function of motor torque are identical in CW and CCW rotating motors. A reduction in load triggers a decline in the number of active stator units in both CW and CCW rotating motors whereas an increase in load promotes an increase in the number of stator units (Fig. 3). In both CCW and CW rotating motors, the kinetics of stator remodeling are determined by the torque exerted by the stator units (Fig. 4). As noted before (29), torque-dependent remodeling in the flagellar motor provides a specific example of how a catch bond (one where binding becomes stronger with tension) can lead to fast mechano-adaptation (34, 35).

The effective free energy difference between bound and unbound units at zero torque (Δ*F*_0_) is positive, suggesting that the binding of stator units is energetically unfavorable. Thus, the production of torque from the motor is required for driving stator assembly. The range of Δ*F* measured here, from around -4 *k*_B_*T* to 6 *k*_B_*T*, is comparable to the energy available from proton motive force (PMF) per proton in a fully energized cell (~ 6 *k*_B_*T*) (36). As torque is proportional to PMF for a given load (36), a lower PMF would restrict the motor to a narrower range of Δ*F* (Fig. 5B) without affecting the slope of the Δ*F* - Γ relationship. Lower torque resulting from a lower PMF would increase *k*_−_ (Fig. 4A) without affecting *k*_+_ in most cases (Fig. 4B), consistent with recent observations in cells affected by the ionophore butanol (30). The values of the model parameters provide quantitative insight into the physio-chemical interactions underlying stator remodeling. The off rate *k*_−_ is the same between the CCW and CW conformations, suggesting that it depends only on the stator-peptidoglycan interactions, which in turn depend on torque but not on the exact mechanisms of torque generation. An increase in the torque exerted by a bound stator unit lowers its effective free energy (Fig. 5B). Torque (Γ) is equal to the radius of the C-ring (*R* ≈ 22.5 nm (5)), multiplied by the tangential force (*f*) applied by MotA on FliG, i.e. Γ = *f R*. Thus, the linear fit in Fig. 5B with slope *λ* = 0.19 leads to: Δ*F* = Δ*F*_0_ – *λ*Γ = Δ*F*_0_ – *fl*, where Δ*F*_0_ ≈ 3.2*k*_*B*_*T*, and *l* is a length scale *l* = *λR* ≈ 4 nm. Assuming the transitions between the bound and unbound state of the stator go through a transition state (Fig. 5A), this length scale denotes the average displacement of the bound state from the transition state along the direction of the force between FliG and MotA, accompanied by a decrease in the free energy equal to the force times the displacement. It is interesting to note that a displacement of 4 nm is within the relevant size range, given that a stator unit is 9 nm high and 7.5 nm wide (15). It is possible that 4 nm represents either an upper limit or a preferred state for the deformation of MotB peptidoglycan binding (PGB) domain.

According to the current (incomplete) understanding, the on rate *k*_+_ of a stator to the motor is controlled by a complex multi-step process. In unbound stator units, the proton channels are blocked by a plug that is a part of the periplasmic domain of MotB (37, 38). Somehow, when an unbound stator unit collides with the rotor, the plug opens, unblocking the proton channel to allow torque generation and the stator unit binds the peptidoglycan (19, 39). First, the interaction between the FliG and MotA drives conformational changes in the stator unit that unplug the channel and extend the periplasmic domain of MotB to the peptidoglycan layer (40, 41). Second, the periplasmic domain binds to the peptidoglycan layer and anchors the stator unit. The first step could be hindered by the rotation of the C-ring, which might impair the interaction between MotA and individual FliG subunits. This could explain the higher on rate during the recovery period, in which the torque is large and motor rotation speed is small. The reason for the independence of *k*_+_ from torque for the low to intermediate torque range (Fig. 4B) remains unclear.

Cryo-EM structures of stator units were obtained recently and provide additional insights into stator function and assembly (15, 16). The structures suggest that CW rotation of the MotA pentamer around the MotB dimer drives rotation of the C-ring. During CCW rotation, the C-ring adopts a narrow conformation and stator units interact with the outside of the ring (32, 42). Upon binding the chemotaxis response regulator CheY-P, the C-ring expands and the stator units now interact with the inside of the C-ring, driving CW rotation (32, 42). These structural changes in the C-ring are the likely source of the asymmetry in the torque-speed relationship of CCW and CW rotating flagellar motors (31, 33). Yet, at the location where MotB binds peptidoglycan, CW and CCW rotation should be indistinguishable because neither the peptidoglycan nor MotB is known to have any CW-CCW asymmetry. This symmetry in the MotB-peptidoglycan interaction is likely responsible for the CW-CCW symmetry in torque dependence of *k*_−_. In CW rotating motors, an unbound stator unit must not only collide with the rotor, but also reach the inside of the C-ring before it can bind the motor. Therefore, the binding of stator units in CW rotating motors is less likely than in CCW rotating motors, consistent with the lower *k*_+_ in CW motors during the recovery period.

Crystal structure of the PGB domain of MotB is known (19). However, the structure provides only limited insight into stator stabilization since it does not reveal the load-dependent deformations in either the MotB PGB domain or the peptidoglycan. Importantly, neither crystal structures nor cryo-tomograms provide information on dynamics. We have shown here and in previous work (28) that motor remodeling depends on dynamic aspects of the stator assembly that are determined by the binding and unbinding of the stator units to the motor. Therefore, a comprehensive understanding of this process requires knowledge of dynamics in addition to the structure. Our work provides this information in quantitative detail.

The biological function of stator remodeling is not fully understood, but both sensory and regulatory roles are possible(10). Load-dependent remodeling of stator units acts like regulated cylinder deactivation in car engines, increasing power output when the demand is high and decreasing output when the demand is low, which serves as a mechanism to save energy. Another likely role of stator remodeling is that of mechanosensitive signaling, in particular, during interaction with surfaces (43–47). Increasing evidence supports the idea that proximity to surfaces increases the load on the motor, which is signalled to downstream processes via a cascade that starts with motor remodeling (48–53). In this manner, stator remodeling might play an important ecological role for bacteria, particularly during the initial stages of biofilm formation.

How cells control the assembly and function of large macromolecular complexes remains a fundamental problem in biology (54–56). It often involves gene regulation, in which a signal of interest triggers a change in the transcription or the translation of genes encoding the assembly components. This process takes at least several minutes and is therefore ill suited for fast-changing environmental signals. The alternative approach, employed by the bacterial flagellar motor, involves direct control of the assembly by the signal of interest. The latter strategy has the advantage that the assembly/disassembly can be triggered directly, allowing the cell to quickly respond and adapt to sudden changes in mechanical cues.

## Materials and Methods

### Bacterial strains and cultures

The strain used in this study (HCB1797; JY32+pWB5+pKF131) was constructed by Junhua Yuan and is previously described (31). Briefly, an in-frame deletion of *fliC* in VS149 [Δ(*cheR-cheZ*)] yielded JY32, which was transformed with two compatible plasmids: pWB5 (AmpR) expressing wild-type *cheY* under an IPTG-inducible promotor, and pKAF131 (CamR) expressing sticky *fliC* under the native promoter of *fliC*. Cells were grown at 33 °C in 10 mL T-broth containing 100 *µ*g/mL ampicillin, 25 *µ*g/mL chloramphenicol, and 0.1 mM IPTG to OD600 between 0.5 and 0.7. Cells were harvested by centrifuging at 1,200 g for 7 min and resuspended in 1 ml electrorotation buffer (20 mM TES, 0.1 mM EDTA, pH 7.5). Flagellar filaments were sheared off by passing the cell suspension through a piece of polyethylene tubing (20 cm long, inner diameter 0.58 mm) 60 times. The cells were pelleted again and resuspended in 5 ml buffer.

### Electrorotation apparatus and data acquisition

The electrorotation apparatus was as described before (28, 57). Briefly, the cells were tethered to a sapphire window in a custom-built flow cell that included the tips of four tungsten microelectrodes a short distance from the surface. Sapphire was used for its high thermal conductivity. The electrodes were driven in quadrature using custom-built electronics. This applied a tunable external torque on the cells tethered on the sapphire surface. The temperature of the sapphire window was sensed by a small thermistor and held at 20 °C by a circular Peltier element driven by a proportional controller. The flow cell together with the electrode assembly was fixed to a 20X objective of a phase contrast microscope. The light diffracted from the cell was split into two parts, one was imaged onto a high-speed sCMOS camera and the other onto a pair of photomultipliers via a linear-graded filter setup (57). The photomultiplier signal was used for live measurement of the motor speed (same as the rotation speed of the cell body) during the experiment and the sCMOS images were used for offline analysis using custom-written MATLAB scripts.

### Data analysis

The data analysis procedure was as described before (28). Angular displacement of the cell between frames was multiplied by the frame rate to obtain the rotation speed, which was filtered by a median filter of order 15. The rotation speed was fitted with steps using a custom algorithm described before (25, 28). The distribution of fitted step heights had two peaks - the first dominant peak due to the addition of a single stator unit, and the second smaller peak due to the addition of two units within a short time interval. We used the unitary step height obtained from the first peak for estimating the number of active stator units from the speed traces.

### Torque-speed (T-S) curve for HCB1797 at 20 °C

The CW T-S curve lacks the characteristic “knee” of the CCW T-S curve, and torque decreases linearly from stall to the zero-torque (31). Additionally, the stall torque and the zero-torque speed of HCB1797 (a derivative of RP437) are smaller than those of the strain HCB986 (a derivative of AW405) that was used in the experiments on CCW rotating motors. The numerical factor for scaling the torque and speed from AW405 to RP437 is 285/350 (31). We therefore derived the T-S curve for HCB1797 by scaling down the stall torque and zero-torque speed for HCB986 (1,260 pN nm and 272 Hz, respectively (28)) and linearly interpolating between those points.

### Hill-Langmuir model for stator assembly

The average kinetics of changes in the number of stator units bound to the motor can be represented by the differential equation,

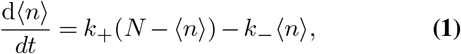

where ⟨*n*⟩ is the ensemble averaged number of stator units bound to the motor at time *t, k*_+_ and *k*_−_ are the on rate and the off rate for the binding of a single stator unit to the motor, and *N* is the number of binding sites, assumed to be 11 (14). The time-dependent solution for an initial condition ⟨*n*⟩ (0)= *n*_0_ is

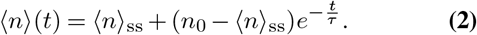

where 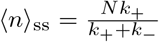 is the steady state number of stator units and 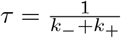 is the time constant for the exponential approach to steady state. We fitted Eq. 2 to the experimentally determined number of stator units, obtaining ⟨*n*⟩ _ss_ and *τ*. Each experimental condition required two separate fits – one for the dissociation of stator units during electro-rotation and another for their assembly after electrorotation was switched off (Fig. 3). From each pair of ⟨*n*⟩ _ss_ and *τ* we calculated *k*_+_ and *k*_−_ for the given value of torque per stator unit Γ specified by the torque-speed curve.

### Model for torque dependence

We developed a model for including torque dependence in the dynamics of stator assembly (28). We assume that at zero torque, the binding of a single stator unit to the motor decreases its free energy by an amount Δ*F*_0_. An increase in motor toque decreases the free energy of a bound stator unit further by an amount *ϵ*_T_ that depends on torque. Thus, the effective free energy difference between the bound and unbound states of a stator unit at a given torque is Δ*F* = Δ*F*_0_ – *ϵ*_T_, where the torque dependence is fully contained in *ϵ*_T_. From equilibrium statistical mechanics, we can get Δ*F* in terms of *k*_−_ and *k*_+_ as 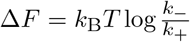, where *k*_B_ and *T* are the Boltzmann constant and the absolute temperature, respectively. We fitted a linear model on the measured value of Δ*F*, given by Δ*F* = Δ*F*_0_ – *λ*Γ, where *λ* is the constant of proportionality. A linear fit on Δ*F* against Γ gave the intercept Δ*F*_0_ as well as the slope *λ*. Assuming that *k*_+_ remains constant, *k*_−_ could be modeled from the linear fit on Δ*F* as 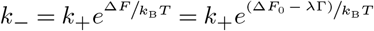.

## ACKNOWLEDGEMENTS

We thank Karen Fahrner and Junhua Yuan for helpful comments on the manuscript. The National Institutes of Health (NIH) supported this work through grants K99GM134124 (to N.W.), R01GM081747 and R35GM131734 (to Y.T.), and R01AI016478 (to H.C.B.).

N.W. and H.C.B conceived the study, N.W. conducted the experiments, N.W. and Y.T. analysed the results. N.W. wrote the manuscript and all authors revised it.

